# Insulin-like peptides regulate oogenesis by stimulating ovarian ecdysteroid production in the Indian malaria mosquito *Anopheles stephensi*

**DOI:** 10.1101/2023.04.06.535964

**Authors:** Benjamin L. Phipps, Mark R. Brown, Michael R. Strand

## Abstract

Females of many mosquito species feed on vertebrate blood to produce eggs, making them effective disease vectors. In the dengue vector *Aedes aegypti*, blood feeding signals the brain to release ovary ecdysteroidogenic hormone (OEH) and insulin-like peptides (ILPs) that trigger ecdysteroid production by the ovaries. These ecdysteroids regulate synthesis of the yolk protein vitellogenin (Vg) that is packaged into eggs. Less is known about the reproductive biology of *Anopheles* mosquitoes, which pose a greater public health threat than *Aedes* spp. because they are competent to transmit mammalian malaria. ILPs can trigger *An. stephensi* ovaries to secrete ecdysteroids. Unlike *Ae. aegypti*, *Anopheles* also transfer ecdysteroids from *Anopheles* males to females during mating. To elucidate the role of OEH and ILPs in *An. stephensi*, we decapitated blood-fed females to ablate the source of these peptides and injected them with each hormone. Yolk deposition into oocytes was abolished in decapitated females and rescued by ILP injection. ILP activity was dependent on blood feeding and little change in triglyceride and glycogen stores was observed in response to blood-feeding, suggesting this species requires nutrients from blood to form eggs. We also measured egg maturation, ecdysteroid titers, and yolk protein expression in mated and virgin females. Although yolk deposition into developing oocytes was significantly reduced in virgins compared to mated females, no differences in ecdysteroid titers or Vg transcript abundance were detected between these groups. 20-hydroxyecdysone (20E) stimulated Vg expression in female fat bodies in primary culture. Given these results, we conclude that ILPs control egg formation by regulating ecdysteroid production in the ovaries.

## Introduction

The perniciousness of *Anopheles* mosquitoes as vectors of human malaria is dependent on the ingestion of blood meals by females for egg production. Of the eight subgenera in the Subfamily Anophelinae (Family Culicidae), only species within the *Anopheles*, *Nyssorhynchus,* and *Cellia* are vectors of *Plasmodium* pathogens (World Health Organization, 2014; Pondeville et al., 2019; Shaw et al., 2022). The two major vectors in the *Cellia* subgenera, *An. gambiae* and *stephensi*, are anautogenous in that females must take blood from humans or other vertebrate hosts to produce a clutch of eggs, and sequential blood meals enable acquisition and transmission of malaria to new hosts. Their reproductive physiology though not as extensively characterized as that of the anautogenous model mosquito, *Aedes aegypti* (Subfamily Culicinae) generally follows the same course (Clements, 1992; Briegel, 2003: Klowden 2007; Shaw et al., 2015). The suite of endocrine factors that regulate metabolic and gonadotropic processes in *Ae. aegypti* is shared with the *Anopheles* females, but little is known about their actions before or after a blood meal, as highlighted below (Zhu and Noriega, 2016; Roy et al., 2016; Strand et al., 2016).

Gonadotrophic cycles in anautogenous females have a previtellogenic phase when primary follicles in the ovaries form and a vitellogenic phase when primary follicles develop into mature eggs in near synchrony. Oogenesis begins in the pupal stage when the 50-70 ovarioles in each of the paired ovaries form primary follicles with an oocyte and seven nurse cells enclosed by follicle cells (Nicholson, 1921; Fiil, 1976; Shalaby, 1971; Laurence, 1977; Clements, 1992; Yamany et al., 2014, 2022). After emerging from the pupal stage, females enter the previtellogenic phase, and during the first two days, juvenile hormone (JH) released from the corpora allata stimulates the primary follicles to approximately double in size before arresting (Hernández-Martínez et al., 2019; Valzania et al., 2019). Females during this phase consume plant fluids and water (Baredo and Gennaro, 2020) to build up metabolic stores for survival and flight until a blood meal is taken from human or vertebrate hosts (Coutinho-Abreu et al., 2021).

Consumption of a blood meal activates the vitellogenic phase by stimulating medial neurosecretory cells in the brain to release neurohormones that directly stimulate gonadotropic processes resulting in egg development, whereas decapitation of *Ae. aegypti*, *An. stephensi*, and *An. albimanus* females shortly thereafter prevents development (Redfern, 1982; Lu and Hagedorn, 1986; Matsumoto et al., 1989). Later work showed such neurosecretory cells in *An. gambiae*, *An. stephensi*, and *Ae. aegypti* contain insulin-like peptides (ILPs) and ovary ecdysteroidogenic hormone (OEH) that restored egg development when injected into decapitated blood fed *Ae. aegypti* females (Brown and Cao, 2001; Wen et al., 2010; Dhara et al, 2013). Five ILP genes have been identified in *An. stephensi* and *gambiae*, and eight ILP genes in *Ae. aegypti*, whereas OEH is encoded by a single gene in the species (Riehle et al., 2002; Krieger et al., 2004; Riehle et al., 2006, Marquez et al., 2011). Whether or not homolog peptides are similarly active in the anophelines has yet to be determined.

In *Ae. aegypti*, ILP3 and OEH both activate insulin/insulin-like growth factor signaling (IIS) in ovaries by binding to closely related receptor tyrosine kinases named the insulin and OEH receptors (IR, OEHR) that are conserved in the anophelines (Brown et al. 2008; Wen et al. 2010; Dhara et al., 2013; Vogel et al. 2015). Activation of IIS in *Ae. aegypti* females stimulates the primary follicles to exit arrest through follicle cell proliferation and production of ecdysteroids that are converted to 20-hydroxyecdysone (20E) (Valzania et al., 2019). Ovaries of *An. gambiae*, *An. stephensi*, and *Ae. aegypti* are similarly responsive to synthetic *An. stephensi* ILP3 and ILP4, thus affiming structural and functional conservation (Nuss and Brown, 2018).

During the vitellogenic phase, ecdysteroid titer peaks 12 - 24 h post blood meal (PBM) and is undetectable by 36 h in different *Anopheles spp.* and *Aedes aegypti* (Hagedorn et al., 1975; Redfern, 1982; Lu and Hagedorn, 1986; Sieglaff et al., 2005; Pondeville et al., 2008 Bai et al., 2010; Pondeville et al., 2008 & 2013; Baldini et al., 2013). In *Ae. aegypti*, 20E signaling along with amino acid sensing/signaling through the target of rapamycin (TOR) kinase pathway and IIS coordinately activate biosynthesis of the yolk protein vitellogenin (Vg) by the fat body for uptake by oocytes in the primary follicles (Roy et al, 2016; Strand et al., 2016). Studies have shown that these signaling elements are engaged during egg development in anophelines (Muema et al., 2017; Hun et al., 2019; Werling et al., 2019; Maharaj et al., 2022), but the extent of their coordinate interactions is not known. 20E treatment stimulates egg development in blood-fed, decapitated *An. stephensi* and *An. albimanus* but not in sugar-fed females (Lu and Hagedorn, 1986; Redfern, 1982, Ekoka et al., 2021), whereas 20E treatment is stimulatory in both blood- and sugar-fed *Ae. aegypti*, a notable difference (Fuchs and Kang, 1981; Deitsch et al., 1995; Gulia-Nuss et al., 2013 & 2015). In *An. stephensi* and *An. gambiae*, amino acid infusion and albumin feeding promoted TOR/IIS and egg development (Uchida et al., 2003; Arsic and Guerin, 2008; Harrison et al., 2021, 2022). Interestingly, treatment of blood-fed *An. gambiae* females with a JH mimic early in this phase depresses ecdysteroid signaling and egg development (Bai et al., 2010). The vitellogenic phase ends by 48 h PBM with chorion covering the yolk filled oocyte, and mature eggs are deposited in a single clutch by ∼72 h PBM. The previtellogenic phase of the second gonadotrophic cycle also begins 48-72 when JH titers increase and secondary follicles in the ovaries develop into primary follicles, which remain arrested until female consumes a second blood meal. (Bai et al., 2010; Hernández-Martínez et al., 2015, 2019; Zhao et al., 2016).

There are two differences in the reproductive physiology of *Anopheles* species and *Ae. aegypti* that warrant further effort to resolve. One is that anopheline adults typically have smaller teneral reserves at emergence and after blood and sugar meals, and depending on nutritional state, may require multiple blood meals for egg development, whereas *Ae. aegypti* typically needs only one meal (Klowden and Briegel, 1994; Timmerman and Briegel, 1999; Telang et al., 2006; Briegel, 2003; Fernandes and Briegel, 2005; Klowden 2007). Other studies of laboratory and wild *An. gambiae* females and other *Anopheles* species, however, do not support this claim (Lounibos et al., 1998; Amed et al., 2001; Oliveira et al., 2012; other refs?). Each gonadotrophic cycle is metabolically costly for females, which rely on nutrients primarily stored as glycogen and neutral lipids such as triacylglycerol (TAG) in the fat body. Most lipids packaged into mature eggs originate from sugars consumed during the previtellogenic phase, while amino acids obtained from the blood meal are used for the synthesis of yolk proteins (Van Handel, 1965; Naksathit et al., 1999a-c; Briegel et al., 2001, 2002; Briegel 2003; Ford and Van Huesden, 1994; Ziegler and Ibrahim, 2001; Wang et al., 2017). Low metabolite stores may underpin the failure of 20E treatment to stimulate egg development in sugar-fed anopheline females (Lu and Hagedorn, 1986; Redfern, 1982). Although the fat body is the principal site for nutrient acquisition and mobilization, studies to date examined nutrient stores in whole bodies and ovaries of anopheline females after sugar and blood meals but not assessed the nutrient dynamics in fat body.

The second difference is that *An. gambiae* and *An. stephensi* males synthesize ecdysteroids in the accessory glands attached to the reproductive tract that are transferred to females during mating (Baldini et al., 2013; Gabrieli et al., 2014; Mitchell et al., 2015; Pondeville et al., 2008, 2019; Bascuñán et al., 2020). Different ecdysteroid forms produced by sex specific enzymes in *An. gambiae* males and females play differential roles in the regulation of receptivity to other males, egg development, and oviposition and that 20E treatment in particular promoted oviposition via c-Jun N-terminal kinase signaling (Peirce et al., 2020; Peng et al., 2022). Ecdysteroids however are not produced in accessory glands of males in the other anopheline subgenera and culicines, including *Ae. aegypti*, and unmated anautogenous females typically complete egg development after a blood meal but fail to oviposit, whereas mated females exhibit monandry after the first mating and deposit eggs. Across these species, unidentified male factors transferred to females block receptivity to other males and promote oviposition (Meuti and Short, 2019). Earlier studies of *An. gambiae* and *An. stephensi* showed that sperm in the spermatheca or a peptide factor activated receptivity and oviposition (Chambers and Klowden, 2001; Klowden, 2001; Ekbote et al., 2003), suggesting that these processes may be regulated by other male factors.

The suite of endocrine factors that regulate metabolic and gonadotropic processes in anophelines is not resolved, and *An. stephensi* is a promising model for such work as demonstrated in earlier studies. With in vivo bioassays, we first addressed whether ILPs, OEH, and 20E have conserved actions in the coordinate activation egg development and vitellogenesis and in blood- and sugar-fed females. The dynamics of nutrient stores in fat body of sugar- and blood-fed females were also profiled to understand whether their state may account for the delay the release of brain factors after blood feeding that activate egg development and the lack of a response by sugar fed females to 20E treatment. We next assessed whether the roles ascribed to male ecdysteroids in female *An. gambiae* are conserved in *An. stephensi* females by comparing egg development and ecdysteroid tissue titers in virgin and mated blood-fed females. The role of 20E in the activation of oviposition by virgin *An. stephensi* females was compared to that of extracts of male and female reproductive tracts to determine if other factors may also modulate this behavior.

## Materials and Methods

### Mosquito rearing

The *An. stephensi* Indian strain originated from the colony at the Walter Reed Army Institute of Research and was reared in an insectary at 28°C and 70% relative humidity on a 12:12 h light/dark photoperiod, as described (Harrison et al., 2022). After hatching, larvae (200/L deionized water) were fed ground TetraMin® fish food daily until pupation, and adults were given a 4% sucrose and 4% fructose solution *ad libidum*. To obtain virgin adults, individual pupae were placed in 1 mL of water in the wells of 96-well plates, and upon eclosion, adults were segregated by sex. Females were fed on a human arm for fecundity and mating experiments or rabbit blood from an artificial membrane feeder for *in vivo* assays with hormone treatment. Afterwards, cotton balls/wicks soaked with sugar solution were provided to groups of females.

### In vivo peptide activity assays

To determine when the brain neurohormones that activate egg development are released after a blood meal, females were decapitated at 0 h and at 6 h increments up to 24 h PBM or left intact. At 48 h PBM, ovaries were explanted to measure yolk deposition. Opaque yolk in three primary follicles of one ovary was measured lengthwise using an ocular micrometer on a dissecting microscope (Leica).

We next tested whether ILPs, OEH, and 20E would restore egg development when injected into blood-fed decapitated *An stephensi.* The bioactivity of synthesized *An. stephensi* ILP3 and ILP4 (>80% purity; CPC Scientific Inc.) used in this study was previously reported (Nuss and Brown, 2018). Recombinant *An. stephensi* OEH (∼18K Da) was expressed in *E. coli* from cDNA (Vectorbase Accession ASTE004312) and extracted and purified by HPLC as described (Vogel *et al*., 2015; Dou et al., 2022). Peptides and 20E (Sigma) were frozen as aliquots of 200 pmol/μl in water and 2 µg/µL, respectively. For hormone treatments, females (3 to 5 days old) were blood-fed, decapitated within 1 h, injected with peptide, 20E, or saline as a control and held, as were intact blood fed females, in plastic cups with access to cotton wicks soaked in water. At 48 h post injection, the number of oocytes with similar amounts of yolk deposition were counted in the ovaries.

### RT-qPCR

To profile *An. stephensi* insulin receptor (*AsIR*; Vectorbase Accession ASTEI02289) expression patterns, abdomens of females 0-4 d post-eclosion (PE) were placed individually in tubes of Trizol reagent. Ovaries (six pairs per tube) and pelts (three per tube) were dissected from females at six-hour intervals 0-72 h PBM. For vitellogenin 1 (*AsVg1*; Vectorbase Accession ASTE003745) expression patterns, pelts (three per tube) were dissected from females at six-hour intervals 0-48 h PBM. After RNA extraction, cDNA templates were synthesized using the iScript cDNA synthesis kit (Bio-Rad). Primers specific for *AsIR* and *AsVg1* were designed and ordered (IDT). Transcript abundance was quantified in reactions containing 3 µL cDNA, 2 µL of 5 µM primers (*AsIR*: forward 5’-CGTAACGGAGTCGGAGAGTG-3’, reverse 5’-CGCCGTTTTCGACTGGATTT-3’; *AsVg1*: forward 5’-TGGTACAACTACACCATCCAGTC-3’, reverse 5’-CTTGATTTCGTTGAGGGTCATGT-3’), and 5 µL of iQ SYBR Green Supermix (Bio-Rad 170-8882) using a Rotor-Gene Q real-time PCR cycler (Qiagen) under the following conditions: denaturation at 95 °C for 10 s and annealing at 60 °C for 45 s, for a total of 30 cycles. An earlier study showed that 20E treatment stimulates egg development in blood fed anophelines but not in sugar fed females. In *Ae. aegypti*, 20E and OEH treatment alone, but not AaILP3 treatment, activate this process in both sugar- and blood-fed decapitated females, thus indicating mobilization of nutrient stores for vitellogenesis by 20E signaling alone or through OEH promotion of IIS, ovary ecdysteroid production, or both (refs). To further explore this differential response in *An. stephensi*, blood-fed decapitated females were injected with 20E or AsILP3 alone or together to assess *AsVg1* expression. Pelts (abdomen body walls with fat body; guts and ovaries removed) were dissected from females at 0, 6, and 12 h PBM, as were pelts from saline injected decapitated and intact females for RNA extraction in Trizol reagent (Ambion) (three per tube). cDNA was synthesized and transcript abundance was quantified as described above. Sugar-fed females (3-5 days old) were injected with 20E or ILP3 alone or together and held as above, as were saline injected and intact blood fed female. At 48 h post injection, the number of oocytes with similar amounts of yolk deposition were counted in the ovaries.

### Triglyceride and glycogen assays

Studies of anophelines have profiled nutrient stores in whole bodies and ovaries of females before and after blood meals but not fat body separately ((Hun et al., 2019, 2021). Stored and blood nutrients are largely taken for egg development, and it is not clear whether store replenishment also occurs in fat body during the previtellogenic phase prior to oviposition. To assess nutrient store dynamics in the fat body of blood fed females, TAG and glycogen levels were quantified in female pelts (five per tube) at 24 h intervals up to 96 h PBM using the Glycogen Assay Kit (Cayman Chemical) and Triglycerides Reagent (Fisher Scientific), respectively.

### In vivo mating assays

*An. stephensi* is in the *Cellia* subgenera as is *An. gambiae*, and evidence suggests males can transfer ecdysteroids to females that may affect egg development and oviposition(Baldini et al., 2013; Gabrieli et al., 2014; Peng et al., 2022; Pondeville et al., 2008). Ecdysteroids were measured in the reproductive tract (RT) and hemolymph of males and females before and after mating at 3 h intervals and days. The last three abdominal segments with the lower RT of males (containing the accessory glands) and females (spermatheca) were placed in 100 µL of 1X phosphate buffered saline (PBS; pH 7.4) (one per tube), briefly sonicated, and centrifuged followed by transfer of the supernatant to a new tube. The rest of the body was crushed in a small chamber with 100 µL of 1X PBS to release the hemolymph, and the solution transferred to a new tube. All samples (6 to 9 replicates for each time) were frozen at -20°C and later thawed to use in an enzyme-linked immunoassay (EIA) to quantify total ecdysteroids (McKinney et al., 2017). Since ecdysteroid signaling plays a role in the activation of vitellogenin gene expression in blood fed females, we assessed whether *AsVg1* expression was altered in mated non-blood fed females in comparison to unmated females. Pelts were collected from virgin and mated females at 1 and 2 days post mating and processed as above to measure expression by RT-qPCR. Ecdysteroid production by ovaries and *AsVG1* expression in fat body were profiled in blood fed virgin and mated females at 0 h and at 6 h intervals during the vitellogenic phase. Two pairs of explanted ovaries with the last three abdominal segments were incubated in 60 µL of saline (N = 3-6 per time) for 6 h at 28°C. Saline samples were frozen, and ecdysteroid content was measured with the EIA (McKinney et al., 2017). Abdomen pelts were collected and processed for RT-qPCR to measure *AsVg1* expression as above.

We next determined whether egg development proceeds similarly in virgin and mated blood fed females of our strain of *An. stephensi*, as shown for a different strain and other anopheline species, including *An. gambiae* (Pondeville et al., 2019; Fig. 4). Sugar-fed virgin females (4 d post eclosion) were placed in a 1×1×1 ft cage with up to 50 virgin males at dusk. After 30 min, several females were dissected to observe spermathecae for presence of sperm and confirm overall mating success. Subsequently, mated and virgin females were blood fed and held separately as above. At 48 h PBM, the number of oocytes with similar amounts of yolk deposition were counted in the ovaries.

Females (3 to 5 days old) with or without access to males were blood fed and virgin females were injected with 20E at 500 ng/0.25 µL or saline as a control and set up similarly. At 72 h PBM, females were placed individually in wells of a 12-well plate with damp filter paper on the bottom to invite oviposition. After 96 h PBM, eggs on the filter paper were counted.

## Results

### ILPs and 20E mobilized after blood-feeding regulate oogenesis

To determine whether the endocrine events that activate the vitellogenic phase are conserved between culicine and anopheline mosquitoes, we first probed whether factors released from the head after blood-feeding stimulate egg formation in *An. stephensi* as occurs in *Ae. aegypti.* To test this, we blood-fed and decapitated *An. stephensi* females and observed yolk deposition into egg chambers 48 h (PBM). Regardless of the timing of decapitation PBM, some females deposited large amounts of yolk while others deposited no yolk, with few females depositing intermediate amounts (Fig. 1). This suggested that *An. stephensi* commits to oogenesis early in the vitellogenic stage and avoids depositing yolk when conditions or resources are unsatisfactory for egg production. However, few females decapitated 12 h or sooner PBM deposited yolk, while the majority decapitated 18 h or later PBM deposited comparable amounts of yolk to intact females (Fig. 1). This indicated that vitellogenic factors were released from the head 12-18 h PBM, and that the timing of this release was later than reported in *Ae. aegypti*. To identify the specific neuropeptides that control egg production in *An. stephensi*, we injected blood-fed, decapitated females with OEH, ILP3, ILP4, or 20E ∼1 h PBM and assessed whether they formed eggs. Both ILP3 and ILP4 fully restored egg production, while OEH and 20E failed to do so (Fig. 2A). Although these results did not support a role for OEH or 20E in anopheline oogenesis, that 20E did not stimulate yolk deposition was surprising given its demonstrated roles in reproduction in *An. gambiae*. Since ecdysteroid titers peak in *An. stephensi* ∼18 h PBM, we next asked whether injecting 20E at a later timepoint PBM to mimic its natural peak can rescue egg formation. To test this, we decapitated blood-fed females 12 h PBM to remove the influence of ILPs, then injected them with 20E, which partially rescued egg production (Fig. 2B). Taken together, these results suggest that ILPs mediate egg production by stimulating ecdysteroid secretion from the ovaries, while 20E mediates egg production by stimulating Vg expression in the fat body.

**Figure 1.**
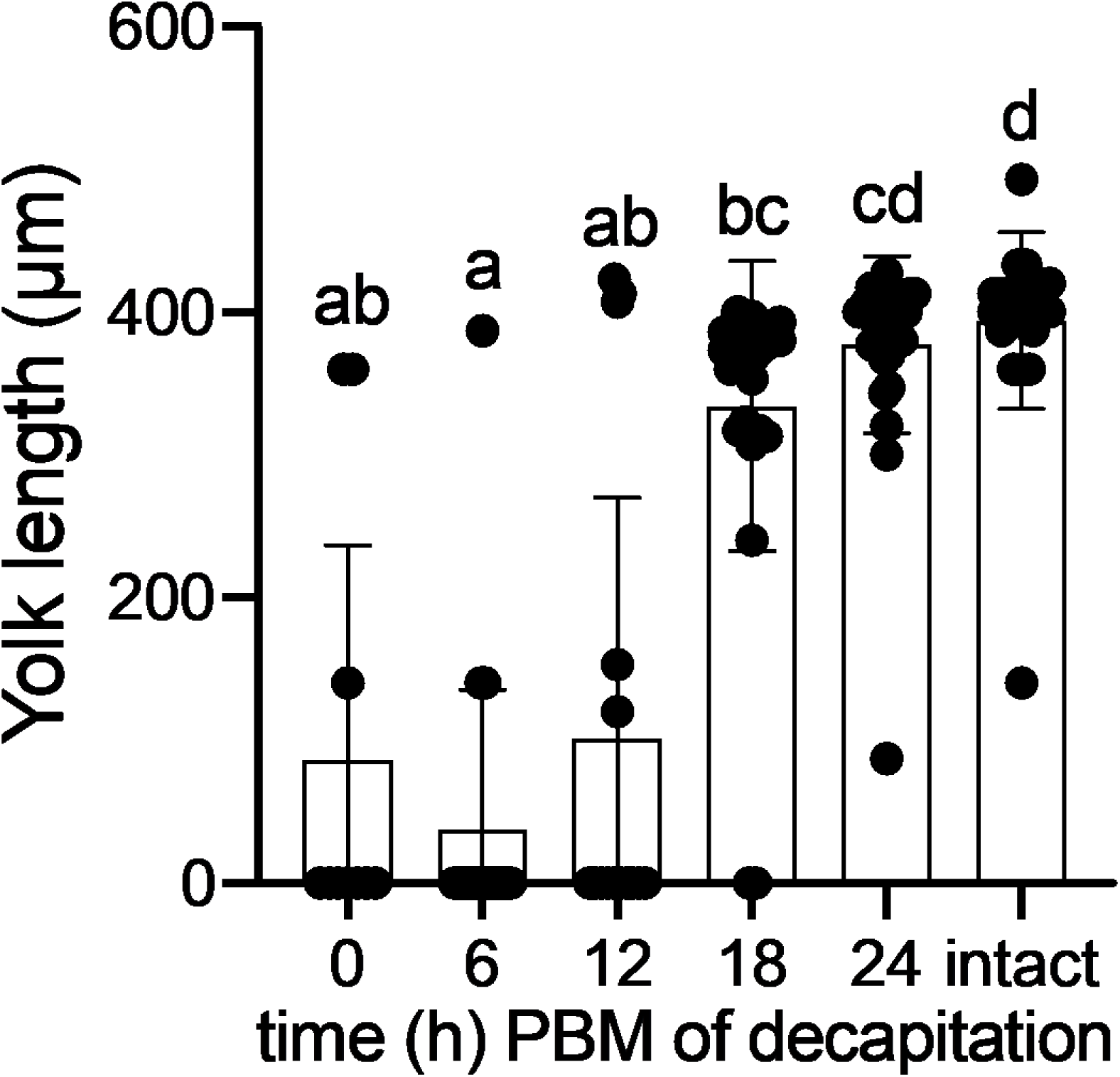
Release of vitellogenic neurohormones from the *An. stephensi* female brain peaks between 12 and 18 h PBM. Number of egg chambers containing yolk (A) and yolk length per egg chamber (B) in females decapitated 0-24 h PBM were analyzed by Kruskal-Wallis test (P = <0.0001).

**Figure 2.**
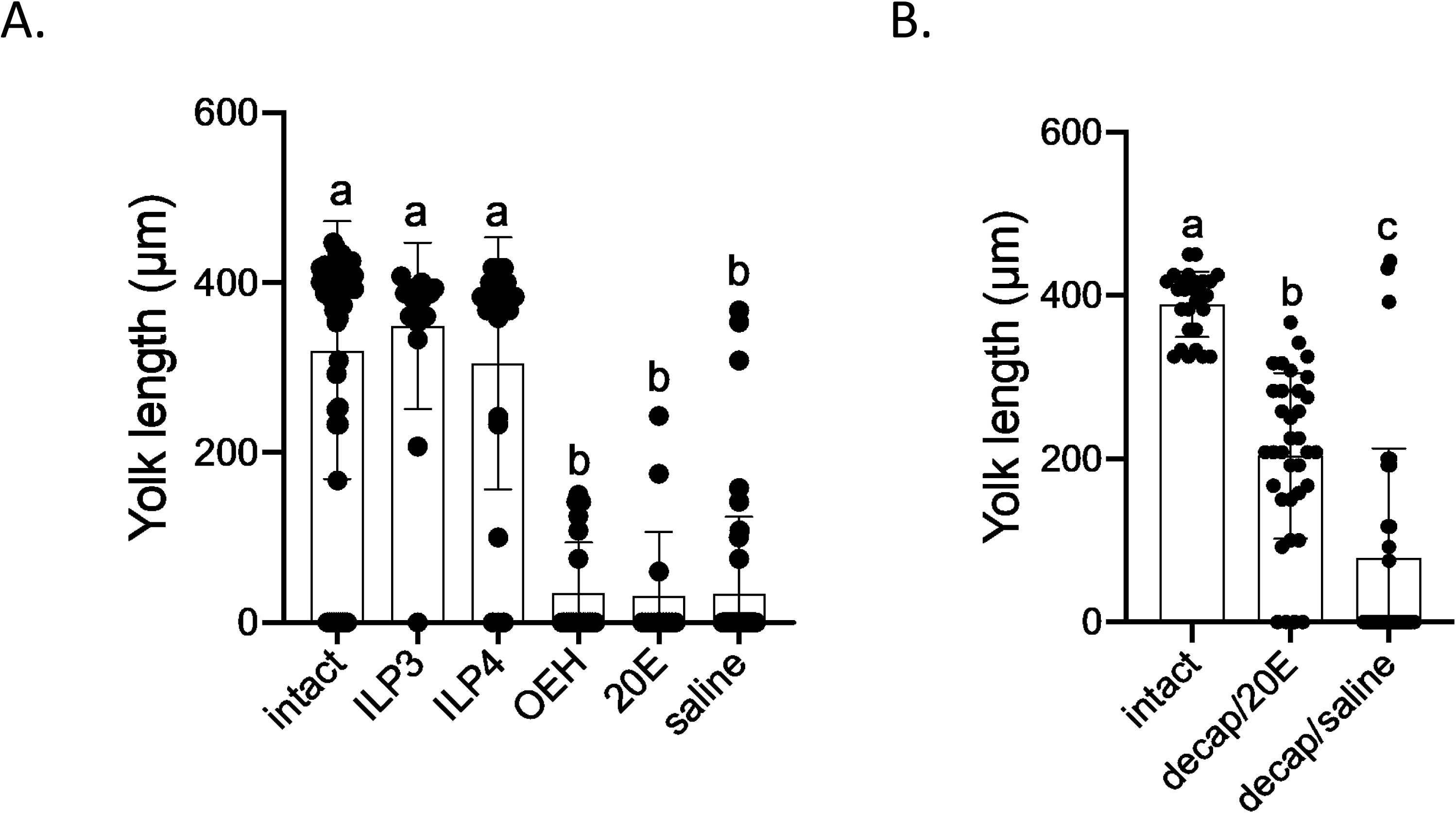
ILPs and 20E stimulate egg formation in blood-fed females. Number of egg chambers containing yolk (A, C) and yolk length per egg chamber (B, D) were compared between blood-fed females decapitated ∼1 h PBM (A, B) or 12 h PBM (C, D) and injected with ILP3, ILP4, 20E, or AS and intact females by Kruskal-Wallis test (P = <0.0001).

In *Ae. aegypti* females, expression of the mosquito insulin receptor (mIR) in the ovaries begins to increase after eclosion and after a few days remains constant during reproductive arrest. Corresponding with development of secondary follicles, mIR expression in ovaries increases again after blood-feeding (Riehle and Brown, 2002). Expression peaks at 36 h PBM and returns to pre-BM levels by 72 h PBM (following oviposition). To compare mIR expression patterns in *An. stephensi* with those of *Ae. aegypti*, we quantified mIR transcript abundance in *An. stephensi* female pelts and ovaries 0-4 d PE and 0-3 d PBM. Like observations in *Ae. aegypti*, mIR expression in *An. stephensi* ovaries and pelts gradually increased for several days PE (Fig. 3A). Another increase was detected beginning at 18 h PBM (Fig. 3B). Expression in ovaries peaked 48 h PBM and returned to pre-BM levels following oviposition (Fig. 3B), while expression in pelts remained unchanged by the blood meal (Fig. 3C). We also quantified Vg expression in *An. stephensi* female pelts 0-48 h PBM. While previous studies report that Vg expression peaks 18 h PBM in *Ae. aegypti*, expression peaked at 24 h PBM in *An. stephensi* (Fig. 4A), further demonstrating delayed activation of endocrine events PBM in this species compared to the dengue vector.

**Figure 3.**
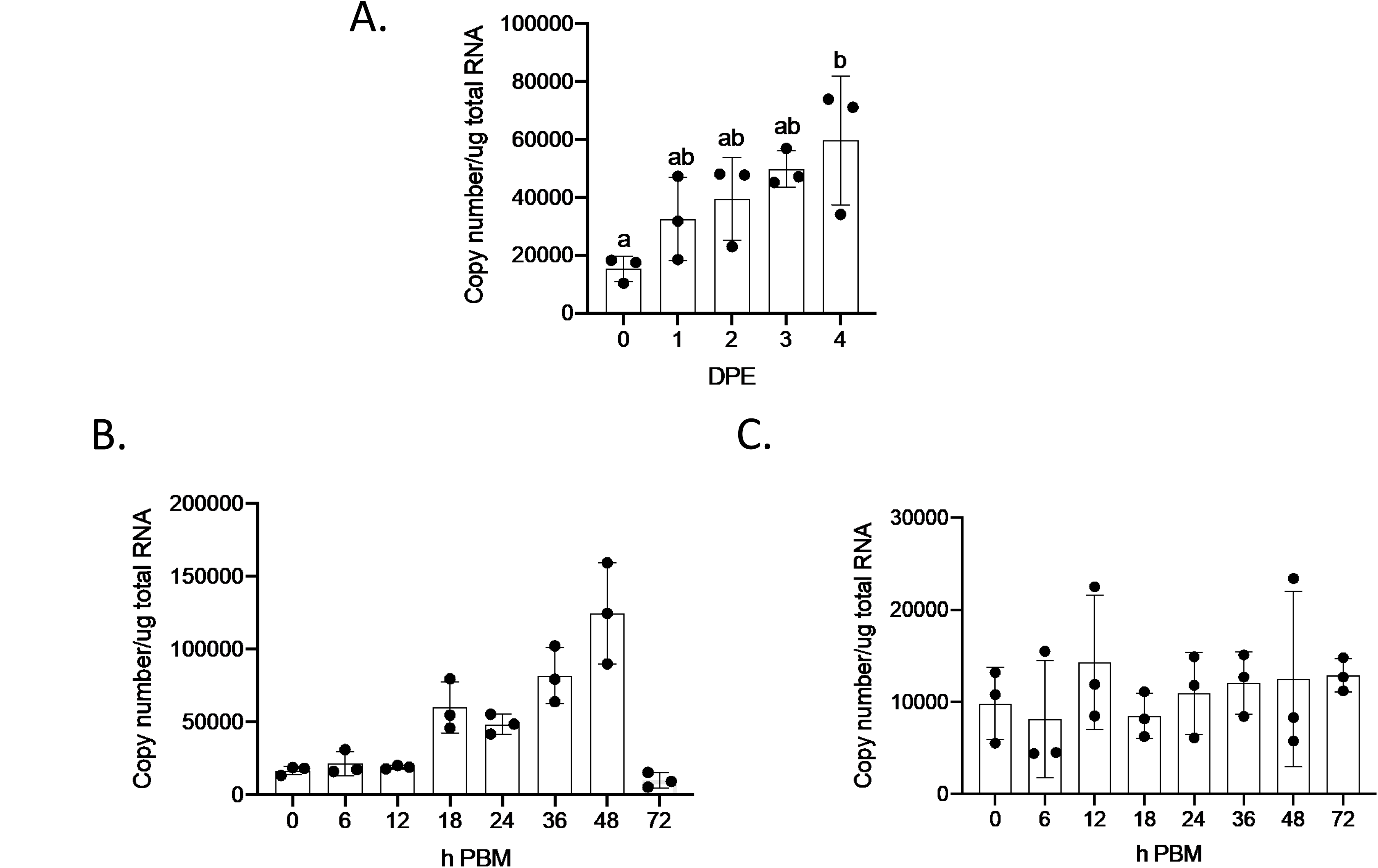
mIR expression increases after eclosion and is upregulated after blood feeding. mIR transcript abundance in abdomens after eclosion (A) and in ovaries (B) and pelts (C) after blood-feeding was quantified by RT-qPCR and results analyzed by one-way ANOVA (A. P = 0.0256, B. P = <0.0001, C. P = 0.2773)

20E and ILP3 synergistically upregulate Vg expression in *Ae. aegypti* female fat bodies (Roy et al., 2007). To test whether this synergism occurs in *An. stephensi*, we injected blood-fed females decapitated at 0, 6, or 12 h PBM with 20E, ILP3, or both. We then measured Vg expression in pelts 24 h PBM using RT-qPCR. In saline-injected females, Vg expression was lowest in those decapitated 0 h PBM and highest in those decapitated 12 h PBM, consistent with gradual release of ILPs from the brain. In 20E-injected females, those decapitated 6 and 12 h but not 0 h PBM demonstrated increased Vg expression compared to saline-injected controls, supporting our observation that 20E can rescue egg formation but not when injected immediately PBM. Finally, ILP3 rescued Vg expression similarly at all time points, and did not further increase Vg expression when injected concurrently with 20E (Fig. 4B). Given that ILP3 has been reported to trigger ecdysteroid secretion by *An. stephensi* ovaries, we concluded that ILP3 activity is localized to the ovaries in this species and does not sensitize the fat body to 20E.

**Figure 4.**
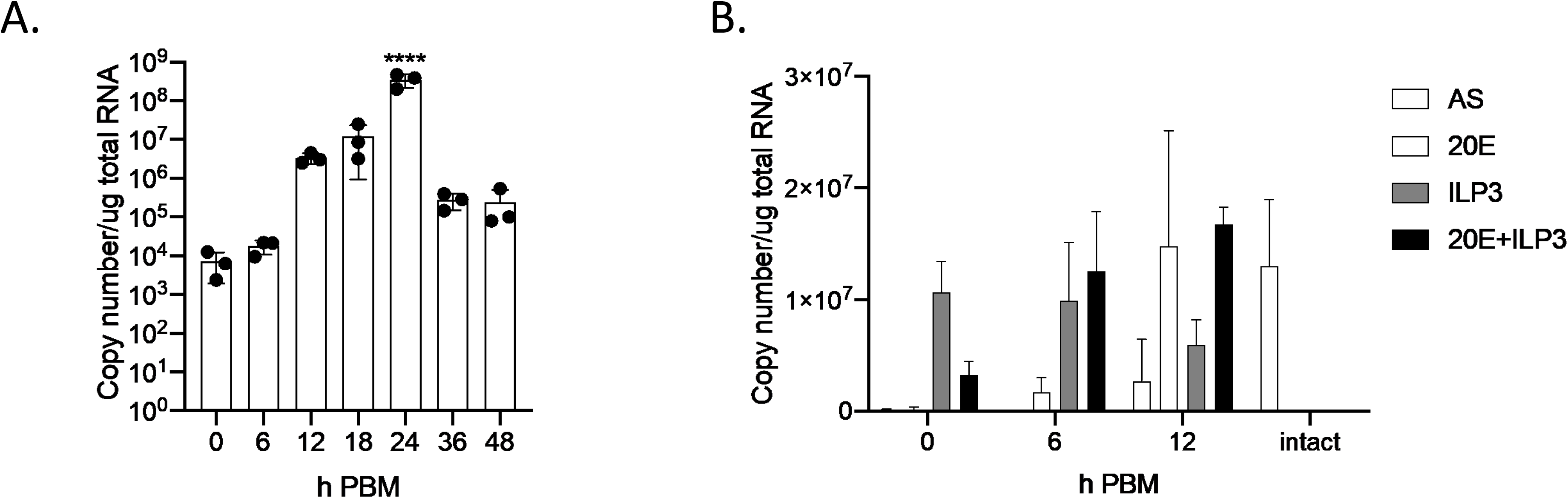
20E and ILP3 upregulate Vg expression but do not synergize in the fat body of *An. stephensi* females. Vg transcript abundance in pelts from blood-fed females 0-48 h PBM (A) and in females decapitated 12 h PBM and injected with 20E, ILP3, both, or neither (B) was quantified by RT-qPCR and analyzed by one-way ANOVA (A. P = <0.0001, B. P = 0.0141).

### Nutrients mobilized after the blood meal are required for 20E-mediated egg formation

Since RT-qPCR revealed that 20E cannot stimulate Vg expression when injected immediately PBM and provided no evidence of a role for ILP3 in sensitizing the fat body to 20E, we reasoned that the factor required for 20E-mediated yolk synthesis likely originates from the blood meal and is released gradually into the hemolymph. To confirm that a blood meal is needed for ILPs and 20E to stimulate egg production, we assessed yolk deposition in sugar-fed, intact females injected with 20E, ILPs, or both. All of these hormones alone or in combination failed to stimulate egg production in the absence of a blood meal (Fig. 5).

**Figure 5.**
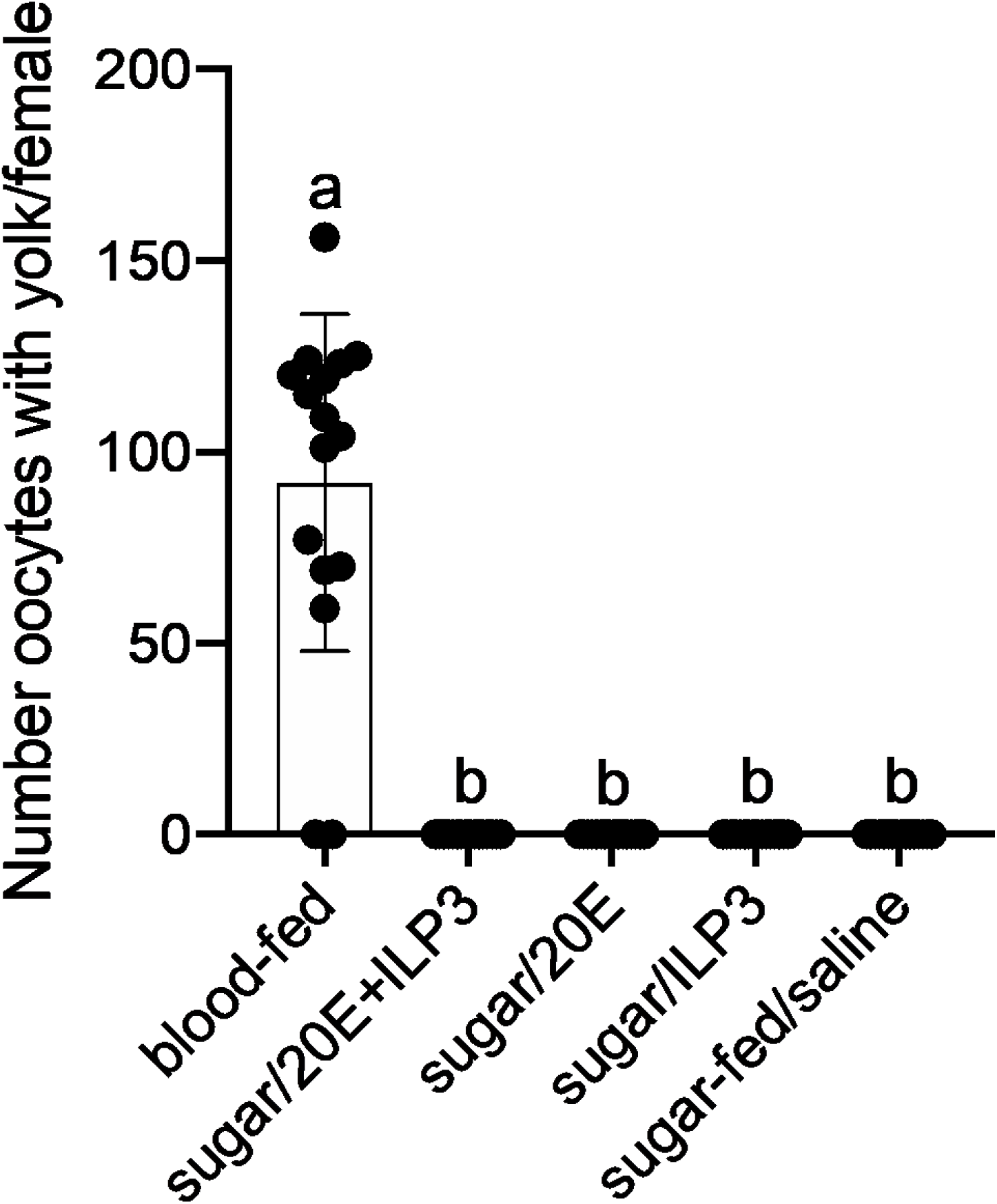
*An. stephensi* females cannot respond to ILPs or 20E in the absence of a blood meal. Number of egg chambers containing yolk (A) and yolk length per egg chamber (B) were compared between sugar-fed, intact females injected with ILP3, 20E, both, or neither and blood-fed females by Kruskal-Wallis test (P = <0.0001).

That *An. stephensi* females were unresponsive to 20E until 12 h PBM was interesting since 20E injection stimulates the mosquito *Aedes aegypti* to deposit yolk into egg chambers even in the absence of a blood meal (Gulia-Nuss et al., 2015; Gwadz and Spielman, 1973). We reasoned that *Ae. aegypti* can use its teneral reserves to produce eggs when stimulated with 20E, while *An. stephensi* must derive nutrients needed for egg formation from the blood meal. After blood feeding, *Ae. aegypti* mobilizes lipids and glycogen stored in the fat body to package into developing eggs (Naksathit et al., 1999). To test whether *An. stephensi* can do the same, we quantified triglycerides and glycogen in *An. stephensi* fat bodies 0-4 d PE and 1-4 d PBM. Females were given the opportunity to oviposit 2-3 d PBM and access to sugar thereafter. Neither triglyceride nor glycogen levels remained unchanged after eclosion and by blood-feeding, oviposition, or access to sugar (Fig. 6), suggesting that *An. stephensi* does not have sufficient teneral reserves to synthesize yolk without a blood meal.

**Figure 6.**
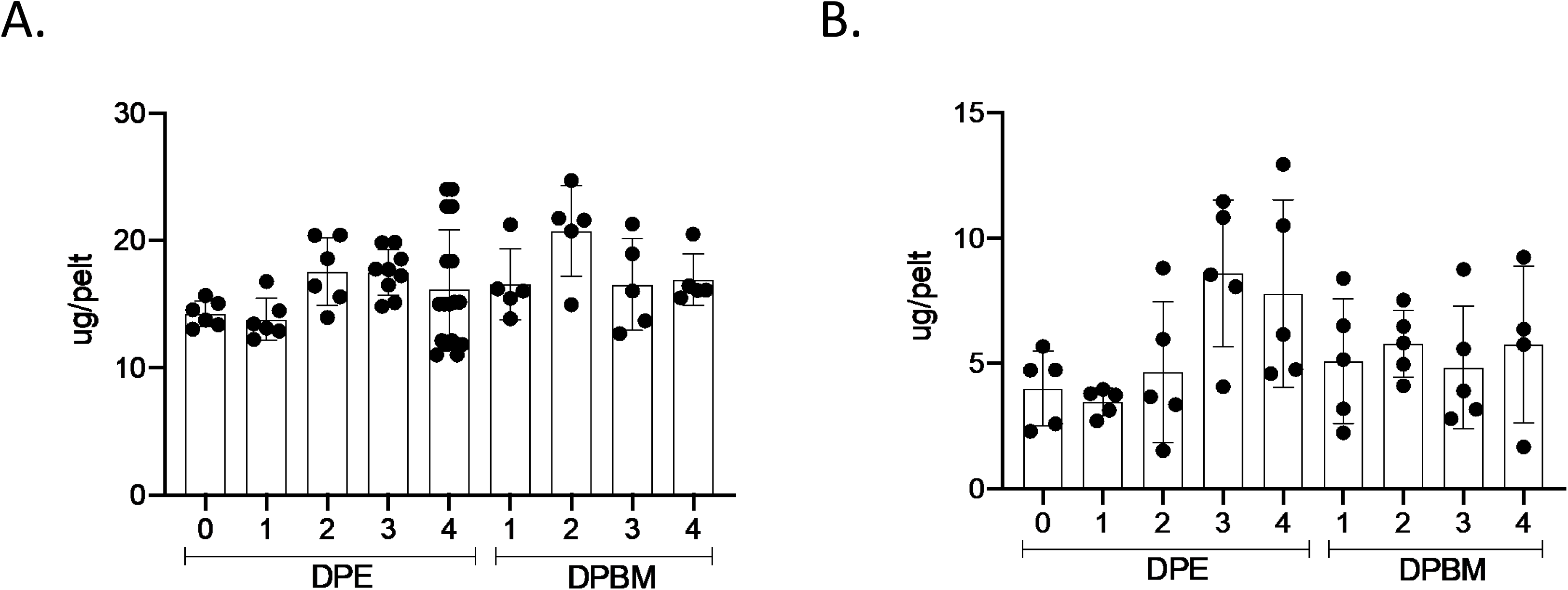
*An. stephensi* females cannot mobilize teneral reserves for egg production PBM. Glycogen (A) and triglyceride (B) levels in the fat bodies of females 0-4 d PBM were measured by colorimetric assay and analyzed by one-way ANOVA (A. P = 0.0004, B. P = 0.0771).

### Mating affects egg maturation but not ecdysteroid titers in females

*Anopheles* mosquitoes in the subgenus *Cellia* can uniquely transfer ecdysteroids from males to females during mating. Since ecdysteroids stimulated the fat body to synthesize Vg (Fig. 4), we reasoned that ecdysteroids transferred during mating could enter the hemolymph and increase fecundity by upregulating Vg expression in females. We tested this using an enzyme-linked immunoassay (EIA) to measure ecdysteroid levels in the hemolymph and reproductive tracts (RTs) of male and female *An. stephensi* before and after mating. RTs but not hemolymph of virgin and mated males demonstrated high levels of ecdysteroids. Freshly mated females had high ecdysteroid levels in their RTs and slightly elevated titers in the hemolymph compared to virgin females, but titers dropped sharply post-mating and were indistinguishable from virgins by 12 h PBM (Fig. 7A). To determine whether transferred ecdysteroids could stimulate Vg expression, we used reverse transcription-quantitative polymerase chain reaction (RT-qPCR) to measure Vg transcript abundance in the fat body before and after mating and found no difference in Vg expression in mated and virgin females (Fig. 7B). These results suggest that ecdysteroids transferred during mating cannot stimulate Vg expression in the fat body.

**Figure 7.**
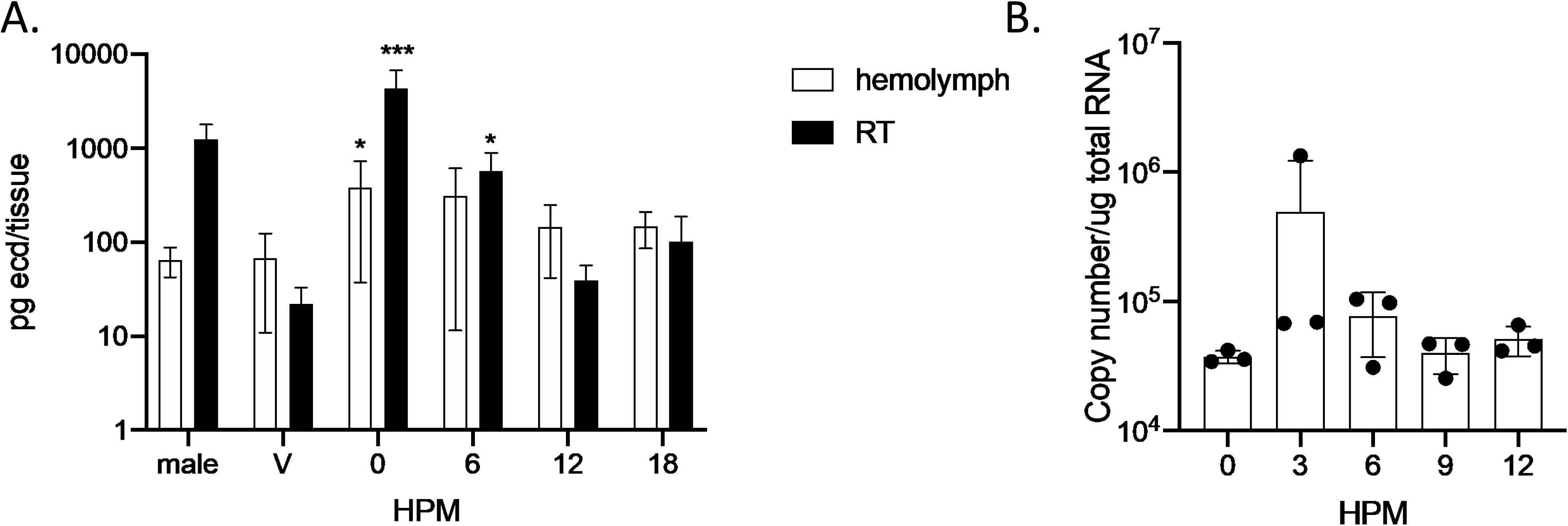
Mating does not influence hemolymph ecdysteroid titers or ecdysteroid secretion from ovaries after mating or blood feeding. (A) Ecdysteroid titers in the hemolymph (white bars) and RT (black bars) in sugar-fed virgin and mated males and in virgin and 0-18 h PM females were measured by EIA and compared by Kruskal-Wallis test (hemolymph P = 0.0277, RT P = <0.0001). Stars indicate significance relative to virgin females in respective tissues. (B) Ecdysteroid secretion by ovaries in sugar-fed virgin and 1-5 DPM females were measured by EIA and compared by Kruskal-Wallis test (P = 0.1964). (C) Ecdysteroid secretion in virgin and mated females 0-48 h PBM was measured by EIA and analyzed by Kruskal-Wallis test (P = 0.1089).

To determine whether mating influences egg production by instead influencing ecdysteroid secretion by ovaries, we explanted ovaries from sugar- and blood-fed virgin and mated females and quantified ecdysteroid secretion by EIA up to 48 h PBM. Consistent with previous results in *An. gambiae* (Pondeville et al., 2008), ecdysteroid titers peaked at 18 h PBM in both mated and virgin females and did not differ significantly between the two groups (Fig. 8A). In agreement with this, mated and virgin females expressed Vg at similar levels PBM, though expression peaked earlier in virgins (18 h) than in mated females (24 h) (Fig. 8B). These results do not support a role for mating in increasing ecdysteroid production by the ovaries or Vg synthesis by the fat body after blood-feeding.

**Figure 8.**
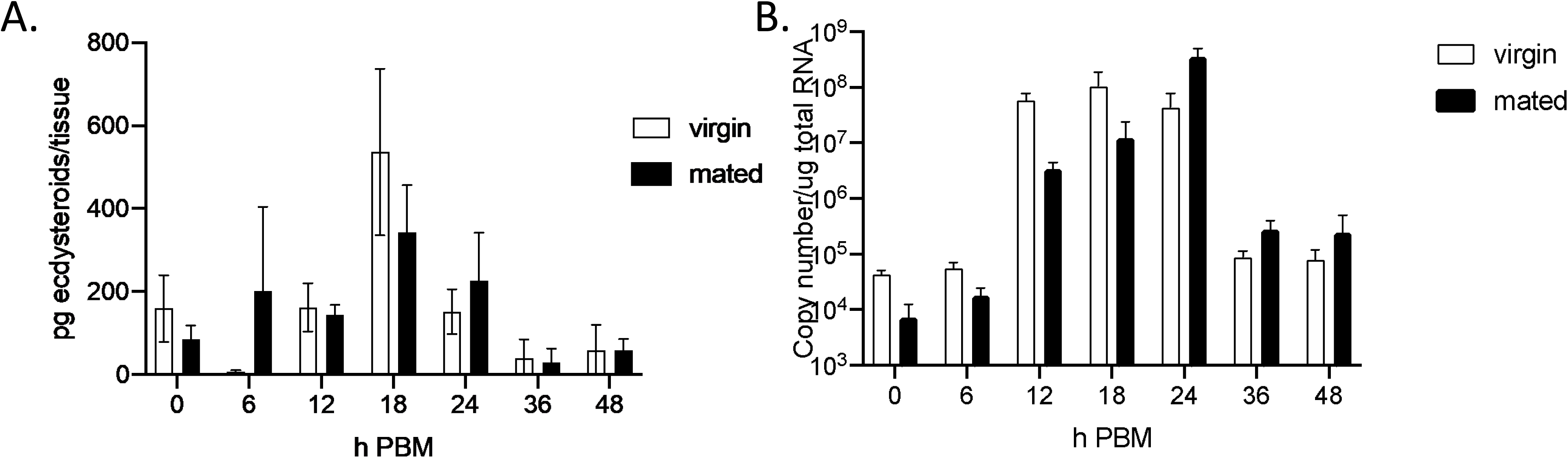
Mating does not influence Vg expression after mating or blood feeding. (A) Differences in Vg transcript abundance in fat bodies of sugar-fed virgin and 1-5 DPM females was measured by RT-qPCR and compared by one-way ANOVA (P = 0.1089). (B) Vg transcript abundance in fat bodies of virgin and mated females 0-48 h PBM was measured by RT-qPCR and compared by Kruskal-Wallis test (P = 0.1484).

To investigate whether ecdysteroids transferred to females during mating influence yolk synthesis, we then probed whether mating influenced fecundity. To do this, we compared egg maturation in mated and virgin *An. stephensi* females after a blood meal by assessing the amount of yolk deposition into egg chambers, as well as the number of eggs matured and oviposited. Like *An. gambiae* (Baldini et al., 2013; Gabrieli et al., 2014), virgin females were less likely to produce yolk after blood-feeding, deposited less yolk in egg chambers, and laid few to no eggs (Fig. 9A). We also tested whether 20E can rescue oviposition in *An. stephensi* by injecting virgin females with either 20E or saline 72 h PBM. 20E failed to rescue oviposition in virgin females, demonstrating no role in reproductive fitness in *An. stephensi* (Fig. 9B).

**Figure 9.**
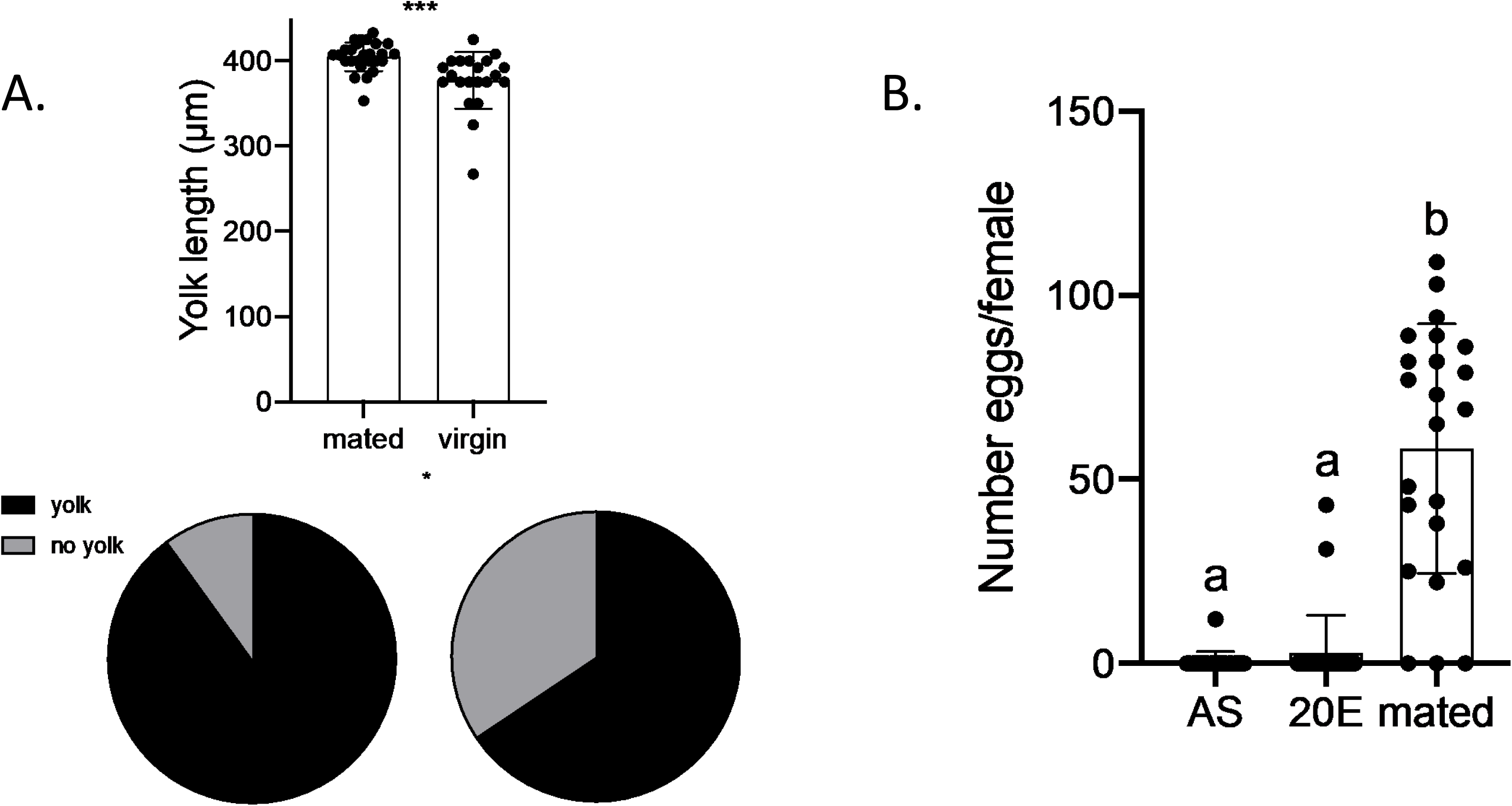
Mating increases yolk deposition in egg chambers and enables oviposition in *An. stephensi* females. Number of egg chambers containing yolk (A), yolk length (µm) per egg chamber (B), and number of eggs laid (C) in blood-fed virgin and mated *An. stephensi* females were compared by Wilcoxon test (A. P = 0.0134, B. <0.0001, C. <0.0001).

## Discussion

Here we provide evidence for conservation of endocrine events activating the vitellogenic phase after blood-feeding among mosquito species. Specifically, we demonstrate in *An. stephensi* that ILPs released from the brain activate ecdysteroid biosynthesis by the ovaries and that 20E stimulates Vg biosynthesis by the fat body. These endocrine functions largely mirror what has been reported in *Ae. aegypti*, though we note several key differences between the species: first, PBM release of vitellogenic neurohormones is delayed in *An. stephensi* compared to *Ae. aegypti*. Correspondingly, the fat body is not immediately sensitive to 20E after blood-feeding. This could be explained by a slower rate of blood meal digestion in *Anopheles* spp.; indeed, aminopeptidase activity peaks earlier in *Ae. aegypti* (24 h PBM) than in *An. stephensi* (30 h) (Billingsley, 1990). Second, like previous studies, we observed an all-or-nothing vitellogenic response to decapitation and hormone injection, suggesting that additional factors inform early commitment to egg formation in *An. stephensi* that are not present in *Ae. aegypti* (Lu and Hagedorn, 1986). Third, OEH stimulates egg formation in *Ae. aegypti* but shows no such activity in *An. stephensi*, possibly indicating that OEH function is divergent in different mosquito taxa. Fourth, we find no evidence of synergism between ILP3 and 20E in stimulating Vg expression in the *An. stephensi* female fat body. Since a prior study demonstrated that ILP3 stimulates ecdysteroidogenesis in ovaries across mosquito taxa (Nuss and Brown, 2018), we conclude that ILP3 elicits a response in ovaries but not the fat body during oogenesis.

Finally, while 20E stimulates egg formation in blood-fed, decapitated *Ae. aegypti* (Gulia-Nuss et al., 2015), no response to 20E was seen in *An. stephensi* unless injected at least 12 h PBM, suggesting that an additional endocrine or nutritional factor released after blood-feeding is required for 20E-mediated yolk synthesis (Redfern, 1982). Furthermore, lipid and glycogen levels in the fat body were low and little change was observed after blood feeding, suggesting that this nutritional input cannot be derived from the teneral reserves. These observations further distinguish this species’ reproductive biology from that of *Ae. aegypti*, which can use its teneral reserves to respond to 20E in the absence of a blood meal (Naksathit et al., 1999). The inability to use teneral reserves for egg production in *An. stephensi* may be explained by life history traits unique to anopheline mosquitoes. Specifically, culicine mosquitoes are better able to build their teneral reserves as larvae compared to anophelines (Timmermann and Briegel, 1993). Therefore, Culicinae mature to adulthood with sufficient nutrient stores to produce eggs, relying on a blood meal primarily to replenish reserves necessary for subsequent gonotrophic cycles (Briegel, 1990a). *Anopheles* spp., on the other hand, lack the teneral reserves upon emergence to make a single clutch of eggs (Briegel, 1990b), and often overcome this problem by taking multiple blood meals to support one gonotrophic cycle (Scott and Takken, 2012). Overall, our results suggest lower nutrient stores and higher dependence on blood meal-derived nutrients in anopheline than aedine mosquitoes.

While we demonstrate transfer of ecdysteroids from *An. stephensi* males to females, change in ecdysteroid titers in female hemolymph after mating is modest compared to those in the RT, and ovarian ecdysteroid secretion and Vg expression remain unchanged. This suggests that ecdysteroids acquired by females during mating may have less impact on diverse tissues than previously thought. We did, however, observe an effect of mating on egg formation and oviposition, consistent with previous results in *An. gambiae*. In that species, mating-transferred ecdysteroids increase fecundity by interacting with MISO in the reproductive tract (Baldini et al., 2013). We conclude that the function of ecdysteroids acquired during mating is distinct from those synthesized after the blood meal and is determined by their localization (female atrium or fat body). Indeed, recent results by other researchers suggest that the female atrium is impermeable to ecdysteroids, preventing dissemination of mating-transferred ecdysteroids via the hemolymph, and that 20E found in the seminal plug has a modification (3D20E) that may distinguish its function from that of 20E secreted by the ovaries. This is logical since if the female were not able to distinguish between the two ecdysteroid sources, mating would upregulate Vg expression, which would be unproductive without a blood meal. Likewise, blood feeding would cause virgin females to lose receptivity to mates, blocking fertilization of synthesized eggs.

## Conclusions

ILPs are released from the brain PBM and demonstrate conserved functions in promoting Vg synthesis by stimulating ovarian ecdysteroid production in both *Ae. aegypti* and *An. stephensi* females. Importantly, ILP mobilization and sensitization of the fat body to 20E are delayed in *An. stephensi* compared to *Ae. aegypti*. This is possibly linked to an increased dependence on nutrients acquired from blood in anopheline compared to culicine mosquitoes. *An. stephensi* females both synthesize ecdysteroids after blood-feeding and acquire them from males during mating, their physiological response depends on the source of these hormones. Specifically, ecdysteroids secreted from the ovaries in response to ILPs act in concert with nutrients from the blood meal to orchestrate Vg synthesis while those transferred during mating have other reproductive functions. Differences in hormone activity among mosquito species should be considered when devising approaches to suppress mosquito populations by targeting reproductive physiology.

## ACKNOWLEDGEMENTS

We thank Jena Johnson and Lilith South for rearing the insects used in this study. We also thank Jai Hoon Eum for synthesizing the OEH homologs used in the *in vivo* assays.

## Author Contributions

B.L.P.: funding acquisition, investigation, formal analysis, validation, writing – original draft; M.R.B.: conceptualization, funding acquisition, formal analysis, resources, supervision, writing – review & editing; M.R.S: conceptualization, funding acquisition, formal analysis, resources, supervision, writing – review & editing.

## Legends

